# From Nets to Molecules: A Comparative Study of Stream Fish Diversity Recovery Using Different Sampling Methods in Eastern Amazonia

**DOI:** 10.1101/2025.01.02.631091

**Authors:** Fabricio dos Anjos Santa Rosa, Cintia Oliveira Carvalho, Anna Karolina Oliveira de Queiroz, Derlan José Ferreira Silva, Silvia Britto Barreto, Birgitte Lisbeth Graae Thorbek, João Bráullio de Luna Sales, Quentin Mauvisseau, Hugo de Boer, Jonathan Stuart Ready

**Affiliations:** Integrated Biological Research Group, Center for Advanced Studies of Biodiversity, Federal University of Pará, Belém, Brazil; Vale Technological Institute, Environmental Genomics, Belém, Brazil; Natural History Museum, University of Oslo, Oslo, Norway

## Abstract

The Neotropical freshwaters of South America host an exceptional level of ichthyofaunal diversity with over 5,160 species, making it the richest continental fauna worldwide. Despite their richness, these freshwater ecosystems face severe threats from human activities, leading to significant declines in fish populations. Traditional fish sampling techniques, such as netting, have been fundamental to ichthyology, offering insights into species richness and abundance. However, the complexity of stream environments limits the effectiveness of conventional sampling tools. As a result, more elusive or niche species are often missed. In recent years, water environmental DNA (eDNA) has emerged as a complementary method to traditional sampling. It allows for detection of aquatic organisms from water samples, expanding the scope of biodiversity assessments. Nevertheless, eDNA filtration faces challenges, especially in turbid waters, including the likelihood of co-extracting inhibitors that can affect amplification and detection processes, as well as the downstream flow of eDNA signals, which means that samples predominantly detect upstream fauna. To address these limitations, the use of bulk samples, such as stomach contents, provides a robust alternative by directly analyzing biological tissues and leveraging the bidirectional mobility of organisms within the stream, enabling the detection of taxa from both upstream and downstream regions. Given these issues, this study combines traditional netting, water eDNA analysis, and dietary metabarcoding to assess the fish biodiversity in three Neotropical streams in the Capim River basin, Pará, Brazil. The integration of multiple sampling techniques offers a more accurate picture of biodiversity, helping to overcome the limitations of each individual method and providing essential insights for conservation efforts.

## Introduction

The South American neotropical inland waters exhibit an extraordinary level of diversity and complexity in their ichthyofauna, hosting more than 5,160 species, which makes it the richest continental fauna worldwide^1^. As a result, this region harbors one in five fish species on the planet and approximately 10% of all living vertebrate species^2^. Despite their remarkable biodiversity, freshwater habitats face disproportionate threats from human activities including climate changes, habitat degradation, forest fragmentation, and pollution^3,4^. Consequently, there is a growing concern regarding the rapid population declines and elevated risk of extinction faced by freshwater organisms^5^. Efficient conservation and management of these unique ecosystems depend on robust and reliable monitoring strategies to assess biodiversity patterns and detect threats early^6^. In this context, accurate monitoring methods are essential to inform conservation policies and to ensure the sustainable use of freshwater resources.

Traditional fish sampling methods, such as netting, have long been the pillar of ichthyological studies, providing reliable data on species richness and abundances^7^. These physical sampling techniques can offer direct evidence of fish presence, allowing researchers to gather critical information about population dynamics^8^ and community structures^9^. However, traditional fish sampling methods face challenges in streams due to the variety of microhabitats, including rocky bottoms, submerged vegetation, and dynamic current velocities^10^, which are often difficult to access. The complexity of these habitats can reduce the efficiency of conventional tools^11^, and even the most versatile net types designed to explore a wide range of microhabitats^12^ are still unable to capture all ecological niches effectively. For example, *Gymnotus* cf. *coropinae* Hoedeman, 1962 was observed foraging at night, always close to the margins, swimming slowly among the submerged roots and trunks, and stalking prey^13^, illustrating how certain species exploit highly specific microhabitats that are challenging to sample with conventional methods. Despite these limitations, traditional methods remain a fundamental tool in biodiversity assessments, serving as a benchmark for emerging techniques like environmental DNA (eDNA)-based monitoring^14^.

In this regard, the eDNA-based approaches were developed with the objective of improving classical microbiology methods to comprehend the variations in microbial diversity within the environment^15^. Its practical applications have rapidly expanded to encompass diverse organisms^16,17^. By using water as an environmental media for eDNA, researchers can now reliably detect and monitor a wide range of aquatic organisms without disturbing them^18,19^. This method marks an improvement over traditional monitoring techniques, which are often hampered by low sampling efficiency and destructive impacts on fish populations^20,21^. Despite its widespread use and advantages, filtration of water faces logistical challenges, particularly in turbid environments with high levels of suspended particulate matter. These conditions not only complicate the filtration process but also increase the likelihood of co-extracting inhibitors, which can hinder downstream molecular analyses such as PCR amplification^22^. Moreover, such conditions can lead to filter clogging after processing only a small volume of water^23^, which in turn may reduce the ability to capture eDNA from rare or low-abundance species and/or later hamper the PCR processes^24^. Other factors, such as acidity, salinity, and temperature have also been proven to affect taxa recovery^25,26^.

In this context, another approach associated with metabarcoding is the use of bulk samples^27^. By utilizing bulk samples from feces^28^ and stomach contents^29^, the metabarcoding approach has proven to be a powerful tool for characterizing the diet of various groups of organisms such as fishes^17^, mammals^30^, birds^31^, and invertebrate communities^32^. One key advantage of this approach is that it directly analyzes tissues, such as partially digested prey in stomach contents or fecal matter^33^, rather than relying on eDNA extracted from environmental samples. Unlike eDNA, which depends on traces of biological material shed into the environment^34^, this method works with physical pieces of the individual, providing a more direct and potentially higher-quality source of DNA. This is particularly important since eDNA is often subject to environmental factors such as temperature, pH, and UV radiation that accelerate DNA degradation, potentially lowering DNA quality and detection rates^35^. However, metabarcoding bulk samples avoids some issues of eDNA but has limitations, including biases related to differential DNA preservation, as some dietary items degrade faster than others, once prey species with hard parts, like exoskeletons, can withstand digestion, potentially complicating DNA extraction for these taxa in laboratory analyses^36,37^.

In general, accurately characterizing community composition from metabarcoding samples is challenging due to limitations in primer selection, which can bias amplification towards certain species, as well as reliance on comprehensive reference databases that may be incomplete for less-studied regions^38^. Likewise, insufficient sequencing depth can limit the detection of rare species, potentially leading to inaccurate biodiversity estimates^39^. Despite these limitations, advances in high-throughput sequencing and primer development are continually improving precision and detection capabilities, making DNA metabarcoding a promising tool for revolutionizing aquatic biodiversity monitoring and conservation efforts^40,41^.

In this study, we aim to describe an integrated monitoring approach for stream fish communities by combining traditional netting techniques with two metabarcoding-based methods: eDNA analysis from water samples and the community fish diets. By leveraging the strengths of each method we seek to provide a more comprehensive assessment of biodiversity. This multi-faceted approach is designed to overcome the limitations of individual methods, offering a more accurate and holistic representation of the biodiversity present in the diverse and complex ecosystems of neotropical streams.

## Methods

### Study Area

The research was carried out in 2022, targeting three first- and second-order streams within the Capim River basin in eastern Amazonia (Fig. 1), with first-order streams being the smallest streams with a clearly defined channel and second-order streams resulting from the confluence of two first-order streams^42^. These small streams were chosen for their marked environmental heterogeneity, making them ideal for exploring different levels of sampling complexity.^43^. Covering an area of 37,000 km^2^, the Capim River basin experiences a humid tropical climate, characterized by distinct rainy seasons from December to May and dry seasons from June to November^44^.

**Fig 1.**
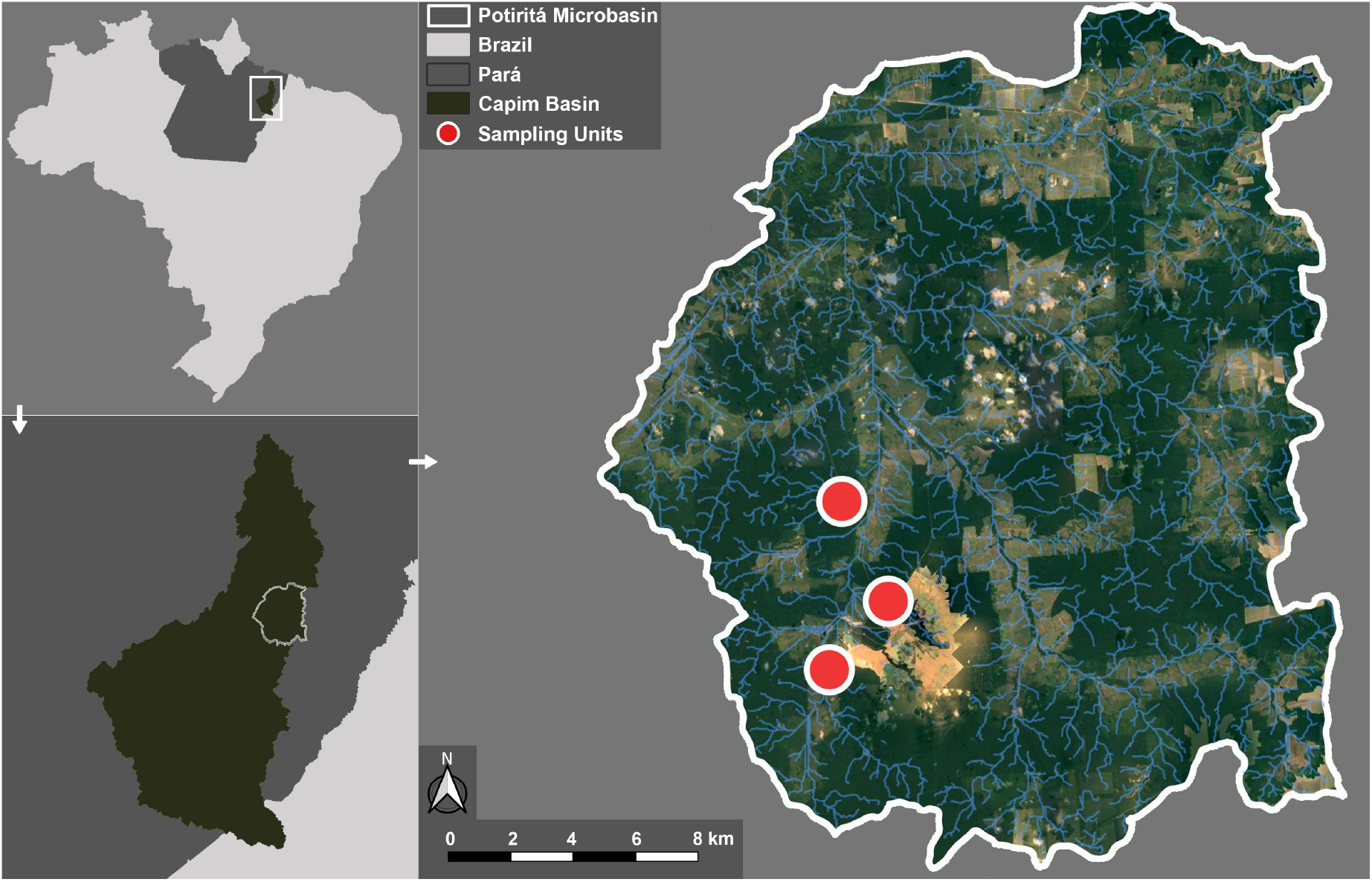
The map illustrates the locations of three sampling units (red dots) in the region of the Capim Basin. The area is situated within the state of Pará, Brazil, with the Potiritá microbasin region outlined in white. The map was generated using QGIS v3.32.0 (https://www.qgis.org). Cartographic parameters included the SIRGAS 2000 Datum and UTM Zone 22S projection, with data sourced from the ESRI Topographic map (30 m resolution), 2024. The microbasin were delineated using the ottobasin coding system^45^, with the classification used corresponding to ottobasin level 5.

### Traditional Sampling

Sampling expeditions were conducted during the dry season to enhance sampling efficiency, facilitated by reduced stream-flow^46,47^, and to minimize variation associated with seasonal environmental changes^48^. To ensure representative samples of species diversity in each stream, the sampling process covered a diverse range of microhabitats, such as litterbanks, sandbanks, and backwater areas. Sampling was conducted using seine nets (width - 2 m; height - 1 m; mesh - 2 mm), hauling 10 times in each stream in a 50-meter section, with each haul spanning approximately two meters across the channel. Samples were preserved in 96% alcohol and identified to the lowest taxonomic level possible using specialized literature and consultation with experts in neotropical fish taxonomy. Collection was conducted under license (SISBIO 68782-5), and all procedures were carried out in accordance with ethical guidelines established by the Federal University of Pará Committee for the Ethical Use of Animals (CEUA 2239161221) following national regulations.

### Water eDNA Sampling

Eight separate water samples, each approximately 400 ml, were filtered from each sampling location, totaling around 3.2 liters. They were distributed across the width of the river being sampled in the beginning (0m), middle (20m) and end of the transect (40m), starting upstream and progressing downstream, with three samples near each bank and three in the center (Fig S1). Water was filtered using a peristaltic pump (Vampire sampler, Bürkle, Germany) and the Swinnex filter holder, using cellulose nitrate membrane filters with a porosity of 0.8 *µ*m and a diameter of 25 mm (Whatman). After filtration, the filters were stored in 1.5 ml Eppendorf tubes with Longmire buffer solution to preserve the DNA until extraction^49^. A field negative control was produced at each location by taking a filter membrane from the packaging and transferring it directly to a tube with Longmire buffer. All tubes were kept on ice during transport and stored in the dark at -20°C until extraction.

### Dietary Sampling

Stomach contents from each fish, obtained through traditional sampling, were removed in the laboratory after using external decontamination protocols (washing unopened stomachs using 5% bleach and then ultrapure water to remove environmental contaminants and loose tissue from the fish) and decontaminated equipment to reduce the contamination risk of the stomach contents^50^. Individual stomach contents were separated from the ethanol preservative by two cycles of centrifugation and washing with ultrapure and UV sterilized water, before being stored whole in separate Eppendorf tubes.

### Molecular Procedures

DNA extraction from filters was performed using the DNeasy PowerSoil kit (QIAGEN) following the manufacturer’s instructions. A negative extraction control was included in each extraction batch. DNA was extracted from stomach samples using an adapted CTAB protocol^51^.

For both water-filtered eDNA and dietary samples, a 313 base pair (bp) portion of the Cytochrome c oxidase subunit I (COXI) gene was targeted using the primer pair mlCOIintF 5’-GGWACWGGWTGAACWGTWTAYCCYCC-3’^27^ and jgHCO2198 5’-TAIACYTCIGGRTGICCRAARAAYCA-3’^52^. This COXI sub-region was selected due to its broad applicability across diverse metazoan taxa and its extensive representation in public reference databases^27^. The primers were synthesized to include doubly unique 12 bp indexes to allow multiplexing of samples in subsequent sequencing procedures^53^.

For aquatic eDNA and the stomach content samples, we used the PCR conditions as follows: 7.5 *µ*L of 2X Accustart Toughmix II (QuantaBio, USA), 5.5 *µ*L of ultrapure H_2_O, 0.5 *µ*L of each indexed primer (10 *µ*M), and 1 *µ*L of extracted eDNA in a final volume of 15 *µ*L. The PCR amplification parameters were as follows: initial denaturation at 94°C for 3 minutes, followed by 5 cycles of 94°C for 10 seconds, 46°C for 20 seconds, and 72°C for 30 seconds, plus 35 cycles of 94°C for 10 seconds, 54°C for 20 seconds, and 72°C for 30 seconds, followed by a final extension at 72°C for 3 minutes. All PCR amplifications were performed in triplicates, along with additional PCR negative control samples.

PCR products were visualized using a 1.5% agarose gel to check for successful production of amplicons. To guarantee equal representation of all amplicons, equivalent quantities of each amplicon were combined using a Biomek4000 liquid handling robot (Beckman Coulter, USA). The amplicons pool were then cleaned using 5% of Illustra ExoProStar (Cytiva, USA) and 0.8X of AMPure XP reagent beads (Beckman Coulter, USA)) and. The amplicon DNA concentration was measured using a Qubit fluorometer with a dsDNA broad-range BR dsDNA BR Assay Kit (Qubit) and following, the amplicon length was confirmed using a Fragment Analyser (Agilent Technologies).

For water eDNA samples, libraries were prepared for the NovaSeq platform (Illumina, USA) with v3 2 × 250 bp kits and spiked with 1% PhiX used as an internal control to calculate error rates (Illumina Inc.) and submitted for sequencing. For dietary samples, libraries were prepared for the MiSeq platform (Illumina, USA) using the v3 paired-end 2 × 250 bp kit and similarly spiked with 1% PhiX to then be subjected to sequencing.

### Bioinformatics and Data Analysis

The quality of sequence data was initially assessed through FastQC reports. The raw paired reads were merged using PEAR 0.9.1^54^, and demultiplexed in OBITools 1.2.13 with the ngsfilter command to separate sequences and assign them to their respective samples based on the unique DNA tags associated with the forward and reverse primers^55^. Sequences longer than 313 bp were removed using the obigrep command. Subsequently, the Obiannotate command removed the specific attributes attached to each sequence read. The USEARCH 11.0.667 software^56^ was used for quality filtering of sequences, dereplication, and clustering, which was performed with a 97% similarity threshold. Based on the previous analyses, tables containing the Operational Taxonomic Units (OTUs) were generated using VSEARCH 2.21.1^57^, serving as the basis for subsequent analyses. Taxonomic assignment was performed using the blastn command^58^ of the Basic Local Alignment Search Tool (BLAST)^59^ against a locally curated reference database. This database was geographically and taxonomically curated based on NCBI data, including all fish taxa registered to occur in Brazil, and was generated using refDBdelimiter tool.

To ensure a more reliable analysis, the maximum number of reads detected in the negative controls for each OTU associated with the sample replicates was subtracted from the corresponding sample values^60^. This step was taken to minimize the impact of false positives, contaminants, or sequencing errors^60^. After subtraction, any resulting negative values were set to zero, and the replicates for each sampling point were summed. OTUs with fewer than ten reads were subsequently removed^14^. Finally, only the species of interest were retained in the final dataset, excluding those exclusively associated with freshwater environments and not occurring in the study area.

For diet samples, the identity of the fish whose stomach contents were analyzed was confirmed by evaluating the taxonomic identities of OTUs with very abundant reads and cross-checked with the previous morphological identification.

To illustrate the distribution of taxa across various sample types, two Venn diagrams were created using the VennDiagram package^61^. The first diagram represents the total number of taxa (OTUs) identified across all taxonomic levels, from kingdom to species, while the second focuses specifically on identifications at the species level. The diagrams highlight the diversity and overlap between sample groups, with unique and shared taxa identified among samples categorized from traditional, water eDNA and dietary sampling. Furthermore, given the differing nature of the three sampling methods, data was analyzed based on frequency of occurrence across the samples for each of the three sample types, producing a heatmap. The heatmap was generated using the package ggplot2^62^. The identified taxa were grouped based on the phylogenetic relationships defined in a well-established phylogenetic tree^63^. All figures were generated using the software R, version 4.4.3^64^.

## Results

The sequencing using a MiSeq platform to analyze fish diets resulted in a total of 5,970,870 reads assigned to bony fishes and rays from 110 stomach samples (∼54,280 reads per sample). In parallel, eDNA analysis from the water samples using a NovaSeq platform generated 14,388,099 reads assigned to bony fishes and rays from 27 (three locations × nine filters) samples (∼532,892 reads per sample). All negative controls showed low read counts overall and no substantial variations between sampling methods, suggesting consistency with minimal background contamination levels, and were therefore removed.

A rich taxonomic diversity of the fish community was identified by all sampling methods (traditional, diet, and water eDNA - Fig. 2). A total of 56 OTUs were identified as bony fishes or rays, including members that represent eight orders, 22 families, and 43 genera. One OTU could only be identified at the family level (Belonidae), while 10 OTUs were identified at the genus level and 45 OTUs were identified at the species level (Fig. 3) Several OTUs were identified, but their taxonomic resolution varies depending on the sample and the identification method used. Some OTUs were assigned to specific species, while others were only classified at the genus level as “*Genus* sp.”, indicating uncertainty about whether they represent unique taxa. These include the pairs/triples: *Astyanax* sp./*Astyanax bimaculatus* (Linnaeus, 1758); *Curimatopsis* sp. Steindachner, 1876/*Curimatopsis crypticus* Vari, 1982; *Pimelodella* sp. Eigenmann & Eigenmann, 1888/*Pimelodella vittata* (Lütken, 1874); *Gymnotus* sp. Linnaeus, 1758/*Gymnotus carapo* Linnaeus, 1758/*Gymnotus coropinae*; *Geophagus* sp. Heckel, 1840/*Geophagus surinamensis* (Bloch, 1791); and *Aequidens* sp. Eigenmann & Bray, 1894/*Aequidens tetramerus* (Heckel, 1840). Other OTUs identified only to the genus level may represent one of many species in the genera *Poptella* Eigenmann, 1908, *Satanoperca* Günther, 1862, *Crenicichla* Heckel, 1840, and *Apistogramma* Regan, 1913, for which multiple reference sequences exist.

**Fig 2.**
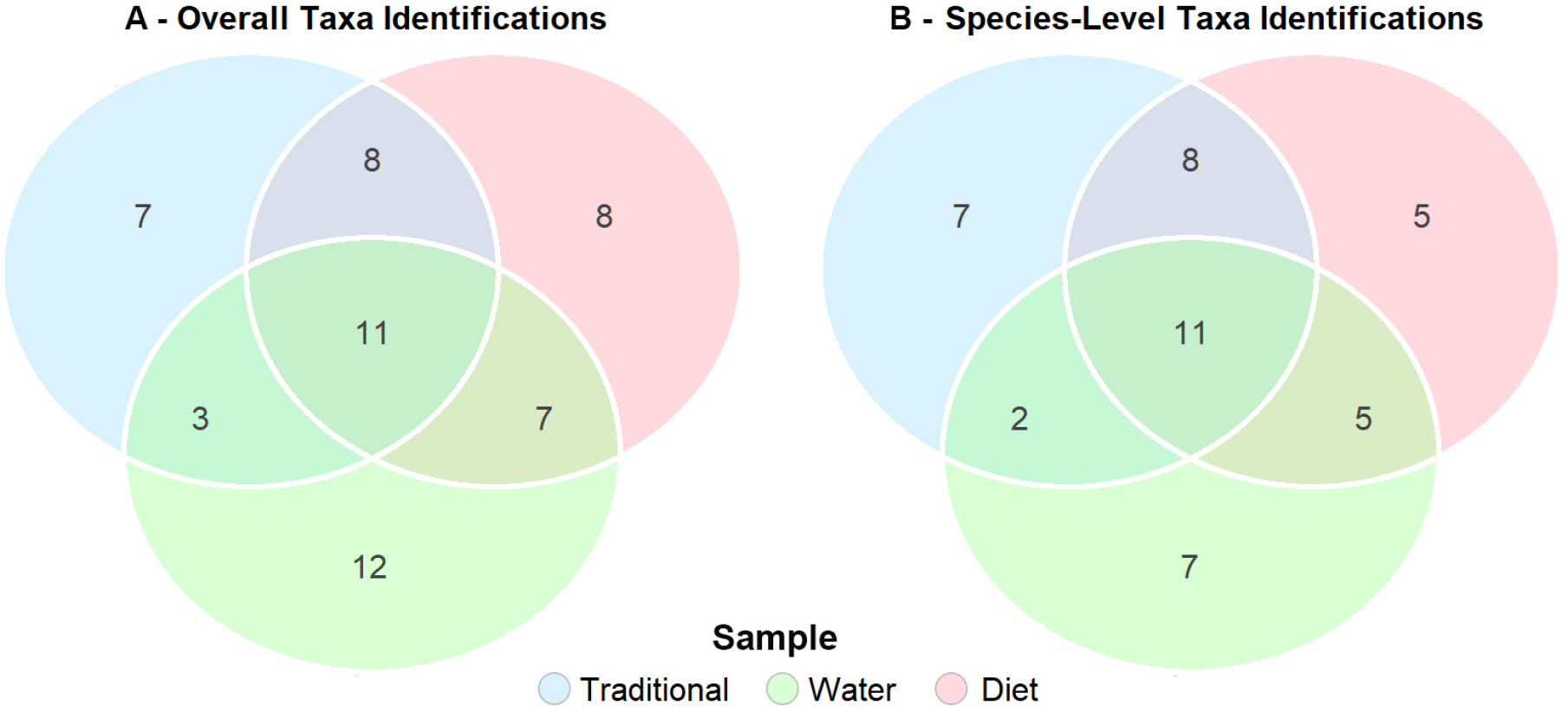
Venn diagram showing the number of taxa detected by each sampling method, highlighting (A) overall taxonomic identifications (family, genus, and species levels) and (B) species-level taxonomic identifications.

**Fig 3.**
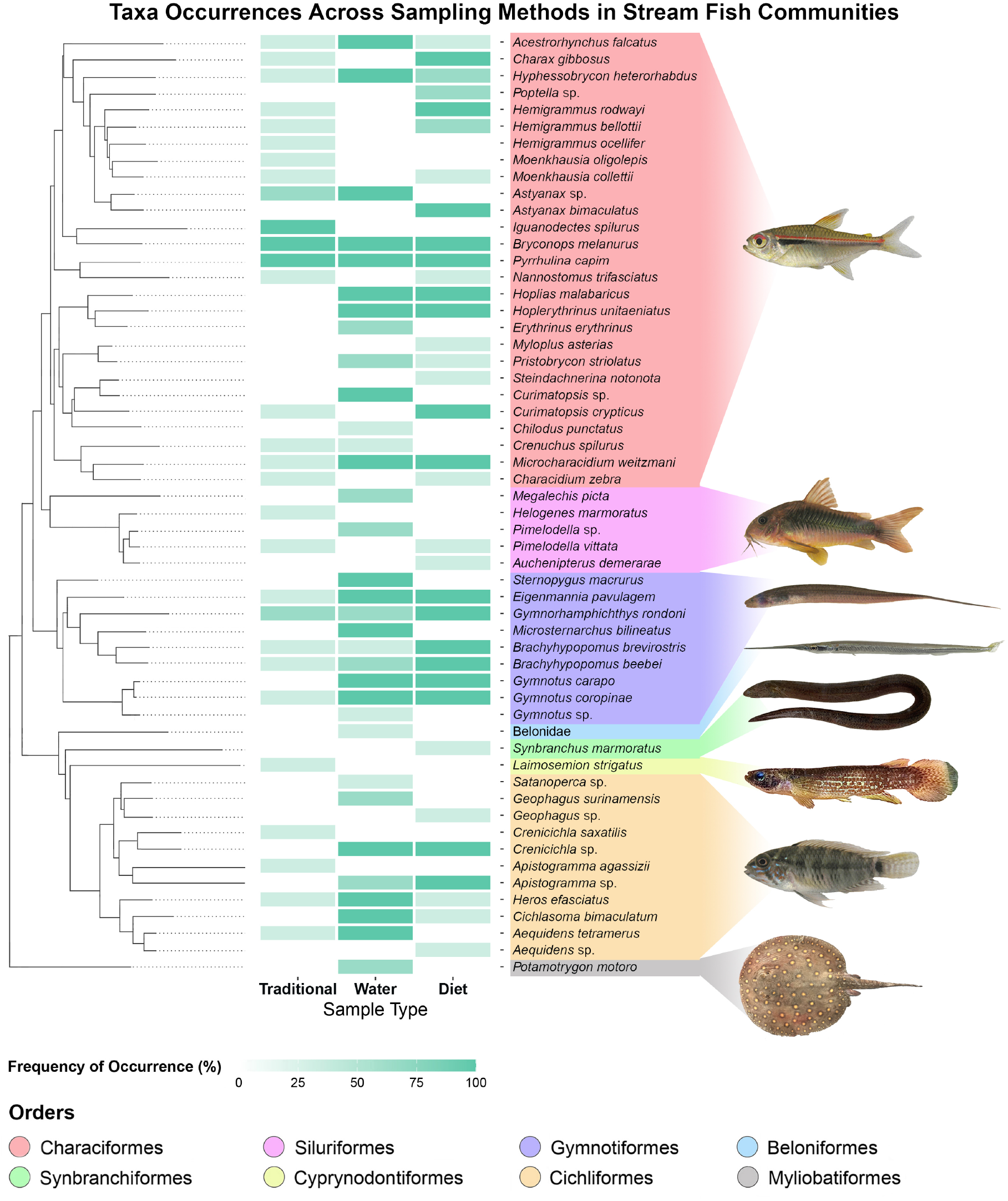
Heatmap highlighting the OTUs grouped by phylogeny with visual representatives of one member for each fish order.

While the raw number of unique taxa identified in Fig. 2 indicates seven unique taxa for traditional sampling, 12 unique taxa for water samples and eight unique taxa for dietary sampling, these numbers are likely marginally inflated due to the differences in taxonomic assignment stated above for OTU pairs or triplets. More careful interpretation of Fig. 3 shows that the traditional sampling approach identified 29 taxa, with one at the genus level and 28 at the species level. Only seven species were unique to traditional sampling (*Hemigrammus ocellifer* (Steindachner, 1882), *Iguanodectes spilurus* (Günther, 1864), *Moenkhausia oligolepis* (Günther, 1864), *Helogenes marmoratus* Günther, 1863, *Laimosemion strigatus* (Regan, 1912), *Crenicichla saxatilis* (Linnaeus, 1758), and *Apistogramma agassizii* (Steindachner, 1875)) (see Supplementary Table S1). Similarly, the water eDNA analysis identified 33 taxa, with one classified at the family level, seven at the genus level, and 25 at the species level (see Supplementary Table S2). Among these, nine species were uniquely detected in the water samples: *Erythrinus erythrinus* (Bloch & Schneider, 1801), *Chilodus punctatus* Müller & Troschel, 1844, *Megalechis picta* (Müller & Troschel, 1849), *Sternopygus macrurus* (Bloch & Schneider, 1801), *Microsternarchus bilineatus* Fernández-Yépez, 1968, *Belonidae, Satanoperca sp*. Günther, 1862, and *Potamotrygon motoro* (Müller & Henle, 1841). Finally, the dietary analysis identified 34 taxa, with five at the genus level and 29 at the species level (see Supplementary Table S3). It included six unique species to stomach content samples (*Poptella sp*., *Hemigrammus bellottii* (Steindachner, 1882), *Myloplus asterias* (Müller & Troschel, 1844), *Steindachnerina notonota* (Miranda Ribeiro, 1937), *Auchenipterus demerarae* Eigenmann, 1912, and *Synbranchus marmoratus* Bloch, 1795).

## Discussion

The exclusive identification of distinct taxa by the different methods shows their complementary value for sampling fish communities (Fig. 2). The ability of each method to recover different components of the ichthyofauna highlights their distinct capacities to access the diverse range of ecological niche spaces occupied by the taxa.

Traditional sampling methods using nets can target a variety of habitats such as backwater pools and lateral ponds^65^. This likely explains why this method was unique in sampling the rivulid *Laimosemion strigatus*, which is typically found in marginal places with low water flow^66^, and where eDNA can quickly settle or degrade without dispersing to the main river body^67^, where we conducted the eDNA sampling. In contrast, traditional sampling using seine nets to catch fish actively can result in difficulties in catching fish from complex habitats with physical obstacles such as submerged roots, or dense vegetation^10^. This is clearly reflected in taxa uniquely identified by both the water eDNA and dietary metabarcoding.

The use of eDNA-based monitoring has previously been shown to recover species that may be elusive, inhabit complex or difficult-to-access environments, or are otherwise less likely to be caught using conventional methods^68,69^. eDNA is also known to be particularly effective in detecting species with cryptic behaviors or that inhabit dense and structured environments, including submerged roots, fallen trees, leaf-litter or sandy/muddy bottoms^13^, such as *Chilodus punctatus, Curimatopsis* sp., *Erythrinus erythrinus*, and *Satanoperca* sp., all of which were found exclusively in our water eDNA samples. Additionally, it is well-suited for identifying species with activity patterns that do not align with traditional diurnal sampling efforts. For example, species that remain hidden in burrows or substrates during the day and are active only during crepuscular or nocturnal periods^13^, such as the catfish and weakly electric knife-fish species identified here (*Gymnotus* sp., *Microsternarchus bilineatus, Pimelodella* sp., and *Sternopygus macrurus*), were also detected exclusively through eDNA water sampling.

Moreover, the use of dietary metabarcoding to assess fish diversity offers some advantages over traditional methods and water eDNA sampling. Using dietary metabarcoding to assess diversity incorporates the natural capability of fish to access microhabitats that are difficult to reach through conventional sampling approaches. Furthermore, the eDNA signal in water follows a downstream flow, meaning that the community detected in any sample predominantly represents upstream fauna^70^. In turn, the diet samples take advantage of the bidirectional mobility of organisms within the stream, enabling the taxa detection from both upstream and downstream regions. Therefore analyzing the diet of fish species becomes a useful approach to identify the presence of rarer or elusive species in various habitats that complement the waterborne eDNA data.

In our study the dietary samples identified nocturnal species such as *Synbranchus marmoratus*, an species that uses the lateral lines to detect and capture prey in absence of light^71^, and *Auchenipterus demerarae*, species that forage alone during the twilight and at night^13^. It can also provide insights into specific interactions that can help us to understand how the diversity is aggregated by this sampling method. For example, the nekto-benthic predator *Gymnotus coropinae* provided evidence of taxa from their environmental space (e.g. *Brachyhypopomus brevirostris* (Steindachner, 1868)), likely ingested in the form of scales or eggs^72^. Otherwise, diurnal channel drift feeders such as *Bryconops melanurus* (Bloch, 1794) and *Moenkhausia collettii* (Steindachner, 1882) were found obtaining signals of small, fast-moving taxa from their environmental space higher in the water column, with evidence suggesting mutual feeding on remnants of each other in the locality where both species are the most abundant members of their trophic guild (see supplementary material S1). Many fishes in Amazonian streams are described as opportunistic feeders, consuming whatever resources are available^73^ depending on the period of the year^74^ and the specific species interactions^75^. However, there are clear stochastic variations in what is detectable through diet that likely represent a combination of behavioral adaptations (predation preferences, scale or mucus eating behaviors^76^), chance opportunities (the eating of tissue fragments released by other predation or other events e.g. sharks in the diet of marine sardines^77^), or the consumption of early life phases^78^.

The complementarity of the three methods likely represents the fact that all of them have their own stochastic variability that affect their respective species accumulation curves in different ways. Traditional fishing methods take a long time to recover rare species that are nocturnal or from difficult-to-sample habitats (e.g. catfishes and weakly-electric knife-fishes) but that do appear in water eDNA and dietary samples. Waterborne eDNA sampling similarly will take more effort to recover species from habitats outside the main water flow that is sampled (e.g. rivulids - *Laimosemion strigatus*), or that release little DNA through natural processes like shedding skin or excreting waste compared to larger species^79^ but that do appear in directed net samples. This may explain the results for some small species like *Iguanodectes spilurus* (Günther, 1864) and *Moenkhausia oligolepis* (Fig. 3). Finally, dietary samples are less likely to provide evidence of apex predators that are not easily caught in nets but that do appear in water eDNA samples such as freshwater stingrays (*Potamotrygon motoro*, Fig. 3) or low abundance species with strong armor such as armor plated catfish (*Megalechis picta*, Fig. 3).

Therefore, combining the analysis of fish diets with their natural mobility patterns provides a more comprehensive approach to identifying multiple fish taxa, especially in habitats where traditional and water eDNA sampling methods may fall short. While water eDNA sampling is a powerful tool, it faces challenges in physically complex stream environments, where disturbances during sampling can lead to false positives or false negatives^80,81^. Additionally, inhibitory substances or sediments selectively collected based on hydrodynamic flow conditions during filtration can hamper PCR amplification, further contributing to false negatives^82^. These challenges, combined with the difficulty of detecting species with specialized habits in such environments, highlight the limitations of both traditional and eDNA methods and emphasize the value of a multi-faceted approach^83^.

In conclusion, our study demonstrates that combining multiple sampling approaches, including traditional methods, diet analysis, and water eDNA, provides a comprehensive understanding of fish biodiversity in complex ecosystems. By integrating these approaches, we can monitor and conserve fish diversity with greater recovery of all taxa, ensuring that management strategies are informed by a holistic view of the ecosystem.

## Data availability

Raw sequencing data can be found here: https://zenodo.org/records/14537156.

## Acknowledgements

We would like to thank Jarl Andreas Anmarkrud and Audun Schrøder-Nielsen from the DNA lab at the Natural History Museum of the University of Oslo for their support and assistance in the lab. We also thank Jessica Conceição Dergan and Marcia da Silva Anjos from Integrated Biological Research Group, who support the team in the field during the data collection trip.

## Author contributions statement

Conceptualization: FASR, JSR; Sampling: FASR, COC, DJFS, SBB; Laboratory analysis: FASR, COC, AKOQ, SBB, BLGT; Bioinformatics: FASR, QM; Data curation: FASR, COC, AKOQ, DJFS, SBB; Statistical analysis: FASR, JSR; Original draft: FASR, QM, JSR; Review and editing: FASR, COC, AKOQ, DJFS, SBB, JBLS, QM, JSR. Funding acquisition: JBLS, JSR, HdB.

